# Transdermal neuromodulation of noradrenergic activity suppresses psychophysiological and biochemical stress responses in humans

**DOI:** 10.1101/015032

**Authors:** William J. Tyler, Alyssa M. Boasso, Hailey M. Mortimore, Rhonda S. Silva, Jonathan D. Charlesworth, Michelle A. Marlin, Kirsten Aebersold, Linh Aven, Daniel Z. Wetmore, Sumon K. Pal

**Affiliations:** Thync, Inc. Boston, MA USA 02199

## Abstract

We engineered a transdermal neuromodulation approach that targets peripheral (cranial and spinal) nerves and utilizes their afferent pathways as signaling conduits to influence brain function. We investigated the effects of this transdermal electrical neurosignaling (TEN) method on sympathetic physiology in human volunteers under different experimental conditions. In all cases, the TEN involved delivering high-frequency pulsed electrical currents to ophthalmic and maxillary divisions of the right trigeminal nerve (V1/V2) and cervical spinal nerve afferents (C2/C3). Under resting conditions when subjects were not challenged or presented with environmental stimuli, TEN significantly suppressed basal sympathetic tone compared to sham as indicated by functional infrared thermography of facial temperatures. In a different experiment conducted under similar resting conditions, subjects treated with TEN reported significantly lower levels of tension and anxiety on the Profile of Mood States scale compared to sham. In a third experiment when subjects were experimentally stressed by a classical fear conditioning paradigm and a series of time-constrained cognitive tasks, TEN produced a significant suppression of heart rate variability, galvanic skin conductance, and salivary α-amylase levels compared to sham. Collectively these observations demonstrate TEN can dampen basal sympathetic tone and attenuate sympathetic activity in response to acute stress induction. Our physiological and biochemical observations are consistent with the hypothesis that TEN modulates noradrenergic signaling to suppress sympathetic activity. We conclude that dampening sympathetic activity in such a manner represents a promising approach to managing daily stress.

## INTRODUCTION

Managing the psychological and physiological effects of stress presents a global challenge for healthcare^1,2^. The negative impact of chronic stress on brain health and cognition has been well defined^3-5^. Unmanaged stress is also known to mediate the breakdown of other biological functions, such as those associated with the immune^6-8^ and cardiovascular systems^9,10^. While there are many effective ways to manage stress and treat stress-related disorders^11^, each approach suffers from some critical limitation for effectively confronting the negative health consequences of stress.

Pharmacological approaches to managing stress, for example with benzodiazepines, suffer from drawbacks like causing drowsiness, blunting general affect, impairing attention and possibly leading to addiction^12^. Relaxation**-**based approaches such as meditation, focused breathing, or mindfulness have been shown to be beneficial, but require time investments and training that often prohibit high compliance rates^13^. Exercise is another effective approach to manage stress and has additional health benefits, but again due to the time commitment required and influence of other motivational factors compliance is often low^14^. To address the above referenced limitations, we hypothesized neuromodulation-based approaches to managing stress might provide a viable alternative.

We sought to develop a method for modulating psychophysiological arousal and stress responses by providing electrical signaling waveforms through afferent pathways of cranial nerves to neuromodulatory nuclei in the brainstem. During signal transmission to cortex, incoming sensory signals carried by the trigeminal and facial nerves simultaneously undergo local processing by a series of highly inter-connected structures including nuclei of the reticular activating system (RAS) located in the pons (Fig. 1a). These brain stem circuits are a first station of information integration in the brain that support higher consciousness by filtering, integrating, and processing incoming sensory information^15^. Here cranial and spinal nerve activity modulates the nucleus of the solitary tract (NTS), locus coeruleus (LC) and other nuclei responsible for the bottom-up regulation of cortical gain, psychophysiological arousal, and neurobiological responses to environmental stimuli and stressors (Fig. 1)^16-22^. More specifically trigeminal afferent activity can modulate noradrenergic neurons via direct projections from the trigeminal sensory nucleus to the LC ^16^. Through these bottom-up pathways, the LC and noradrenergic signaling modulates human behavior, sleep/wake/arousal cycles, and higher cortical functions including attention and cognition ^16-22^. Exemplifying functional modulation of this circuitry, trigeminal nerve modulation has been shown to be moderately effective for treating psychiatric disorders like depression ^23^ and neurological conditions like epilepsy ^24,25^.

**Figure 1.**
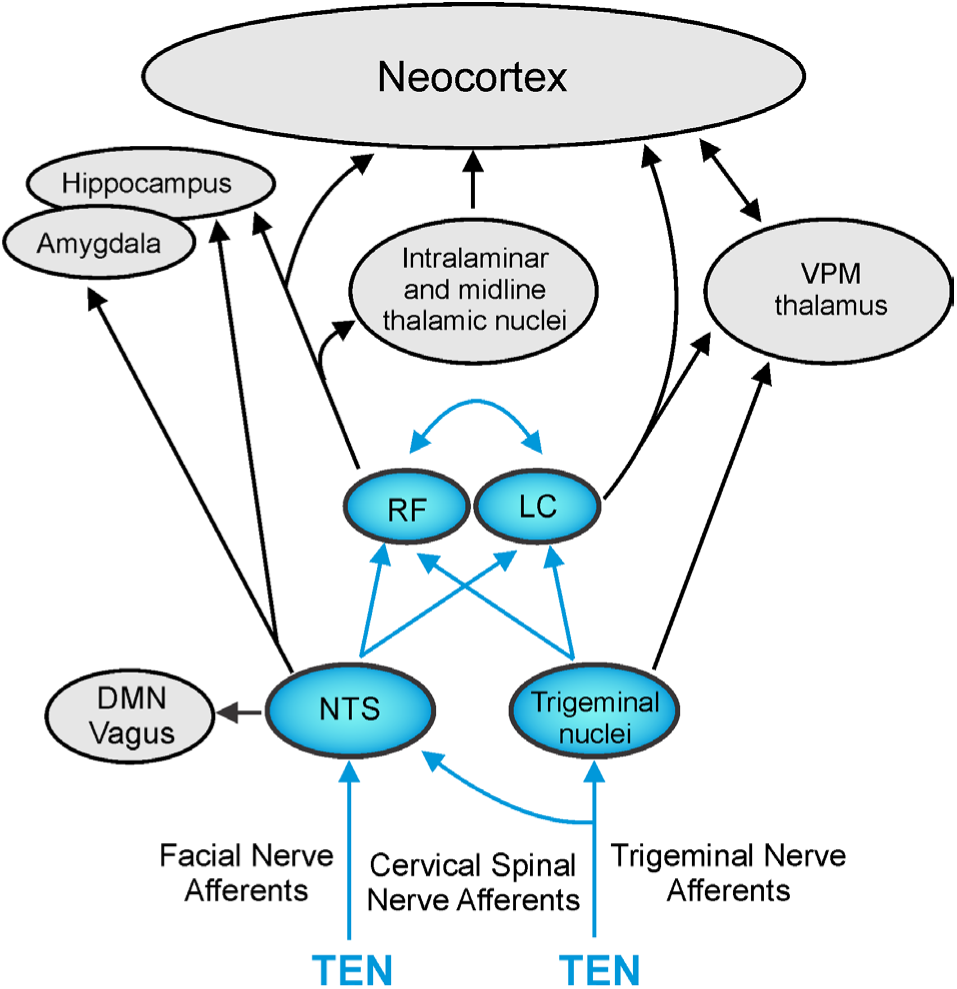
Cranial and spinal afferent pathways provide direct inputs to neural circuits regulating psychophysiological arousal and stress responses. The diagram illustrates trigeminal and facial nerve afferents impinging on the nucleus of the solitary tract (NTS) and trigeminal nuclei (primary sensory nucleus and spinal nucleus) in the brainstem. The NTS and trigeminal nuclei impinge on the noradrenergic locus coeruleus (LC) and other brainstem nuclei comprising a loose network of interconnected structures historically recognized as the reticular formation (RF). The major pathways and circuitry highlighted (blue) were targeted using high-frequency pulsed currents (7 – 11 kHz) using transdermal electrical neurosignaling (TEN) waveforms delivered to trigeminal, facial, and cervical spinal (C2/C3) afferent fibers. Trigeminal and facial nerve afferents (sensory and proprioceptive) can modulate the activity of the LC and NTS to affect psychophysiolgical arousal, the activity of the sympathoadrenal medullary (SAM) axis, biochemical stress responses, and the dorsal motor nucleus (DMN) of the vagus. The bottom-up neurosignaling pathways illustrated serve mechanisms for the regulation of cortical gain control and performance optimization by influencing a broad variety of neurophysiological and cognitive functions, as well as primary sympathetic responses to environmental stress^26,27^.

Given the growing interest in electrical and electromagnetic neuromodulation methods, we have spent the past few years developing wearable neurosignaling systems, devices, and approaches that enable the safe and comfortable modulation of peripheral and central neural pathways using pulsed electrical waveforms. Norepinephrine (NE) mobilized from the LC plays an integral role in mediating the brain’s major sympathetic responses to environmental stressors, making it a promising target for mitigating stress responses. In the present paper, we tested the hypothesis that neuromodulation employing a high-frequency, pulse-modulated transdermal electrical neurosignaling (TEN) waveform targeting afferents (sensory and proprioceptive) of the right trigeminal nerve (V1/V2 distributions), temporal branch of the facial nerve, and cervical spinal nerves (Fig. 2a) can dampen basal sympathetic tone and alter the psychophysiological and biochemical responses to experimentally induced acute stress. Our observations suggest this neurosignaling approach may be useful for alleviating the psychological, physiological, and biochemical responses to acute stress by exerting an effect on endogenous noradrenergic signaling.

**Figure 2.**
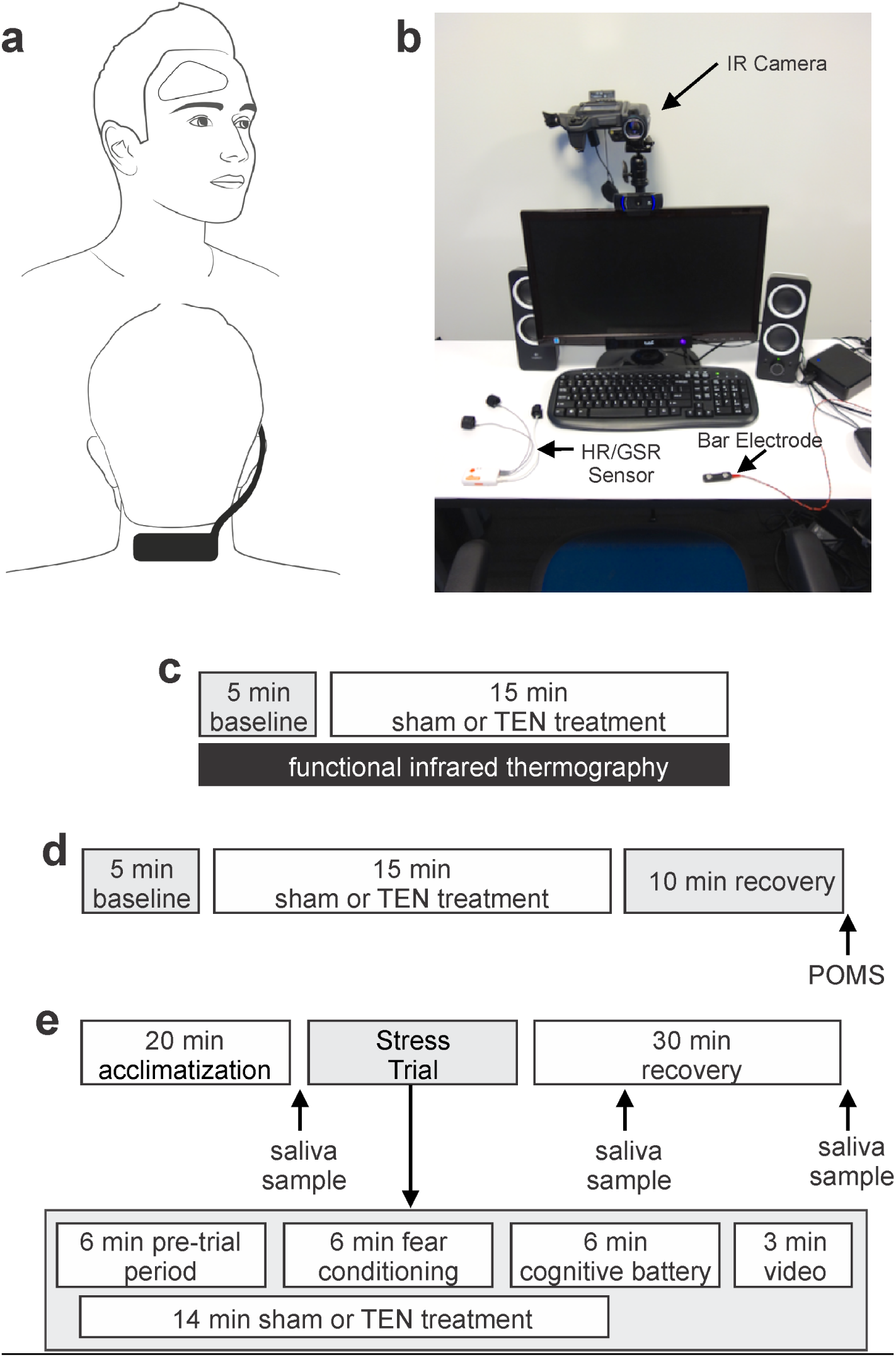
Experimental approaches used to assess the effects of TEN on basal sympathetic tone, mood, and acute stress responses. **a**, The photographs illustrate the montage and device used to target cranial and cervical spinal afferents. **b**, The photograph illustrates the testing room and shows an infrared (IR) camera, integrated heart rate and galvanic skin conductance sensors, a bar electrode used to deliver fear conditioning stimuli, and a monitor for presenting stimuli. In sub-panels **c** – **e** schematics illustrating the three experimental paradigms used in this study are shown. **c**, The schematic depicts Experiment 1 designed to evaluate the influence of TEN and sham stimulation on a sympathetic skin response by monitoring emotional thermoregulation of facial temperatures using functional IR thermography. **d**, The schematic depicts Experiment 2 designed to investigate the influence of TEN and sham stimulation on affective mood states. **e**, The schematic depicts the experimental approach used to evaluate the influence of TEN and sham stimulation on psychophysiological and biochemical responses to stress. For additional details, see *Methods*.

## RESULTS

### The modulation of sympathetic skin responses by TEN as indicated by functional infrared thermography

In Experiment 1, to begin evaluating the influence of TEN on sympathetic activity, we monitored a sympathetic skin response (SSR) using functional infrared thermography of facial regions (Figs. 2c and 3). Thermoregulation of skin temperature is influenced by physiological homeostasis, as well as changes in autonomic nervous system activity imparted by emotional responses to stimuli^28,29^. The SSR (sudomotor activity) includes rapid changes in skin conductance and skin temperature due to micro-perspiration, muscle activity, and vasodynamics (blood vessel constriction/dilation)^28^. Functional infrared thermography has shown that increased sympathetic activity leads to the rapid temperature decreases of the nasal and mouth regions of non-human primates^30,31^, as well as several facial regions of humans^32-34^. Conversely decreased sympathetic activity, increased positive emotional states or relaxation lead to temperature increases of several facial regions of humans including the forehead, nose, cheeks, and mouth region including the chin^35-37^.

**Figure 3.**
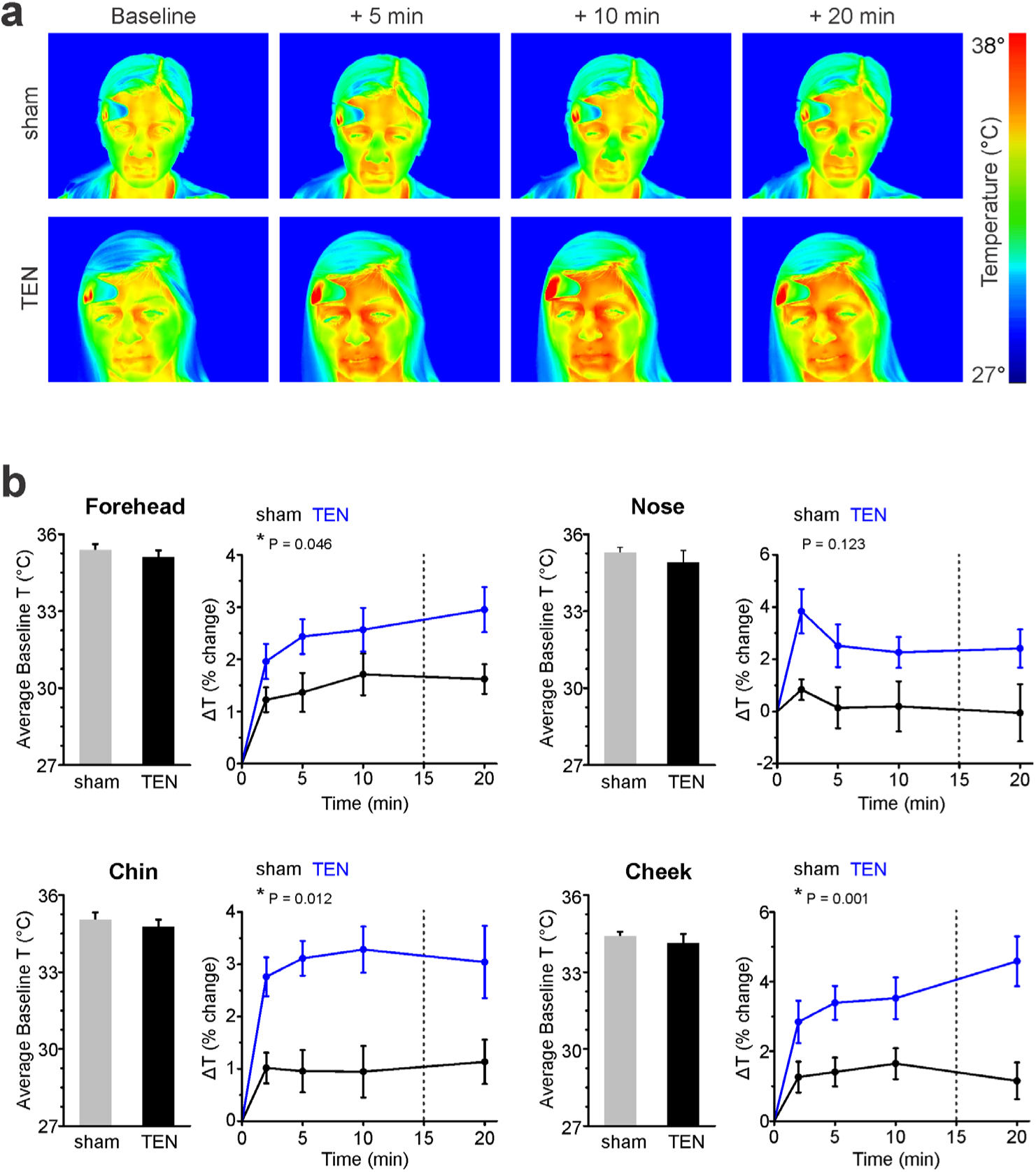
TEN significantly modulates a sympathetic skin response and emotional thermoregulation by increasing facial temperatures as indicated by functional infrared thermography. **a**, Pseudo-colored infrared (IR) images are shown for representative subjects from the sham (top) and TEN (bottom) treatment groups obtained in Experiment 1. The average baseline temperature and time points corresponding to 5, 10, and 20 min following the onset of a 15 min sham or TEN treatment are illustrated. **b**, Histograms (left) illustrate the average baseline temperatures obtained from the foreheads, noses, chins and cheeks (Supplementary Figure 1) of subjects during the 5 min baseline period prior to sham or TEN treatment. The lineplots (right) illustrate the average percentage change in temperature (ΔT) of foreheads, noses, chins and cheeks of subjects at time points corresponding to 2^nd^, 5^th^, 10^th^, and 20^th^ min period after the onset of sham (black) or TEN (blue) treatment (Supplementary Videos 1 and 2). The vertical dahsed lines in the line plots illustrates the termination of sham or TEN treatments. The P values for groups differences calculated from repeated measures analysis ANOVA’s are shown and an asterisk indicates a significant group difference at P < 0.05. For clarity all data have been shown as mean ± SEM.

Using a between subjects design, we monitored temperature changes of the forehead, nose, cheeks and chin in response to sham (n = 10) or TEN treatment (n = 9; Supplementary Figure 1). Briefly, volunteer subjects were randomly assigned to one of the two treatment groups and blinded to the experimental condition. At the beginning of the trial, thermographic recordings were acquired during a 5 min baseline period, during the 15 min sham or TEN treatment period, and during a recovery period after treatment (Fig. 2c). There were no significant differences in the average temperatures of the facial regions between subjects during the five-minute baseline period (Fig. 3b). A two-way repeated measures ANOVA with time as the within subject factor and treatment condition (sham versus TEN) as the between subjects factor was conducted on each facial region monitored: forehead, chin, cheek and nose. Both sham and TEN produced a significant increase in the temperature change from baseline for all facial regions (forehead: F(4, 68) = 46.66, P = 8.25E-19; chin: F(4, 64) = 17.39, P = 1.0E-6; cheek: F(4, 68) = 29.00, P = 4.13E-9; nose: F(4, 68) = 4.47, P = 0.016). This main effect was subsumed by significant treatment-by-time interactions. Compared to sham, the temperature increases were greater for TEN across the times measured for the forehead, chin, and cheek (forehead: F(4, 68) = 4.73, P = 0.046; chin: F(4, 64) = 4.61, P = 0.012; cheek: F(4, 68) = 8.26, P = 0.001; Fig. 3b and Supplementary Videos 1-2). There was not a significant treatment-by-time interaction for nose temperatures based on treatment condition (F(4, 68) = 2.18, P = 0.123). These data indicate that TEN can significantly dampen basal sympathetic tone compared to sham in a manner sufficient to modulate emotional thermoregulation as reflected in temperature changes of the face.

### The influence of TEN on transient affective mood states

In Experiment 2 (Fig. 2d) we implemented between subjects design using a different group of volunteers to determine the influence of sham (n = 20) and TEN (n = 25) on transient mood states using self-reported responses on the Profile of Mood States (POMS) questionnaire^38^. The POMS is a validated and widely implemented scale designed to measure fluctuating or transient moods across six affective states or subscales: Anger-Hostility, Depression-Dejection, Fatigue-Inertia, Vigor-Activity, Tension-Anxiety, and Confusion-Bewilderment. As described above volunteer subjects were seated in a testing booth (Fig. 1b) where they rested quietly for a five-minute baseline period before receiving a 15 min sham or TEN treatment. Ten minutes following the completion of their treatment, subjects completed the 65-item POMS survey (Fig. 2d).

We performed a series of one-way ANOVA’s to test for group differences between TEN and sham stimulation on each POMS subscale. Participants who received the TEN treatment reported significantly less anxiety and tension (POMS score = 1.73 ± 0.52) compared to those who received sham stimulation (POMS score 2.10 ± 0.64; F(1, 44) = 4.51, P = 0.04; Fig. 4). There were no discernable group differences on the remaining mood subscales: Anger-Hostility: F(1, 44) = 1.96, P = 0.169; Depression-Dejection: F(1, 44) = 3.11, P = 0.085; Fatigue-Inertia: F(1, 44) = 0.206, P = 0.652, Vigor-Activity: F(1, 44) = 0.19, P = 0.667, and Confusion-Bewilderment: F(1, 44) = 0.26, P = 0.613. These data demonstrate that TEN significantly reduced self-reported tension and anxiety compared to sham when subjects are not challenged or presented other environmental stimuli.

**Figure 4.**
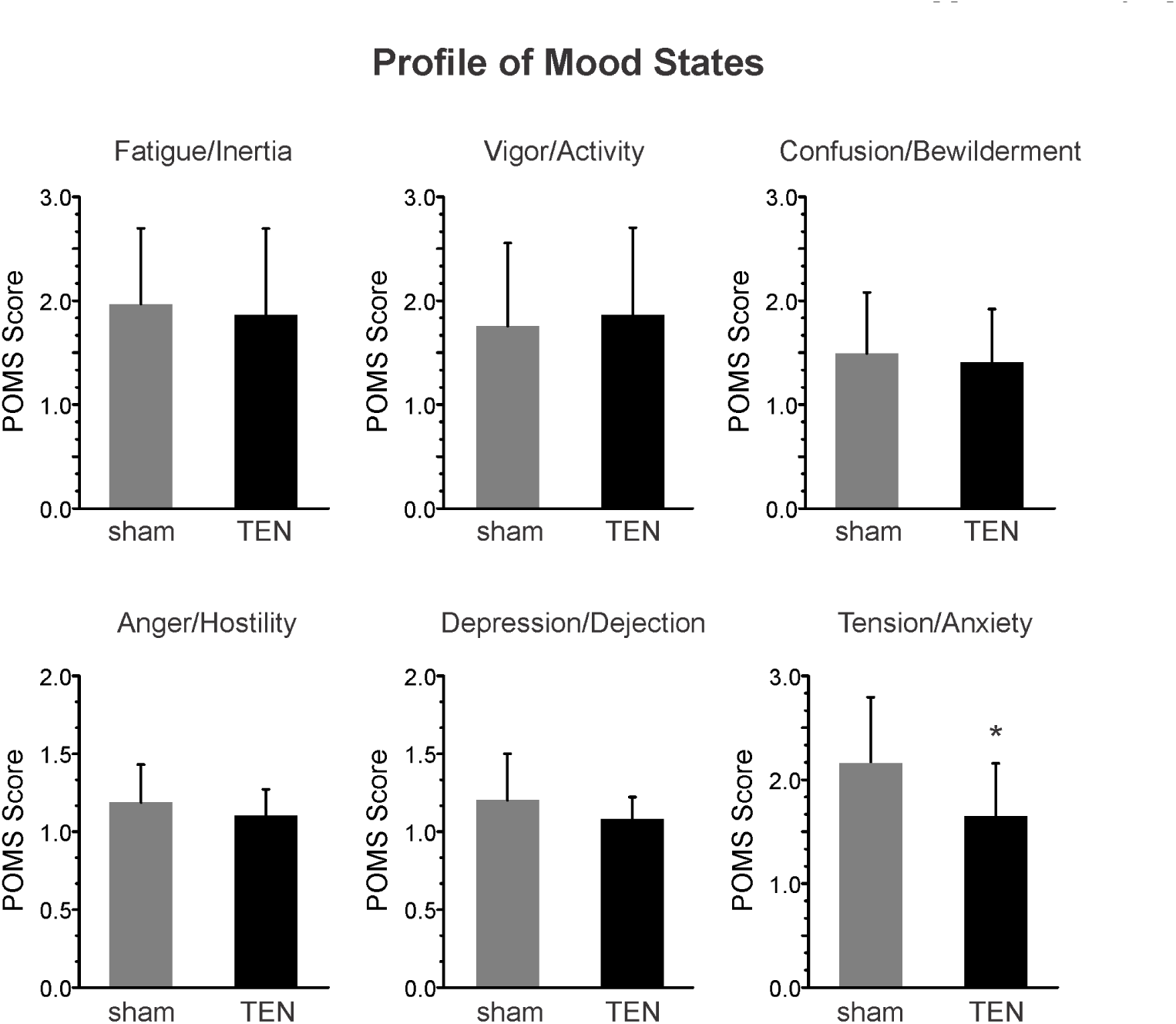
TEN significantly reduces anxiety and tension compared to sham. Data obtained from Experiment 2 are illustrated. The histograms illustrate the mean ± SD scores obtained from TEN and sham treatment groups on the Profile of Mood State (POMS) subscales: Fatigue/Inertia, Vigor/Activity, Confusion/Bewilderment, Anger/Hostility, Depression/Dejection, and Anxiety/Tension. Following a 15 min treatment session, subjects in the TEN group reported significantly less Anxiety/Tension on the POMS survey compared to the sham treatment group. An asterisk indicates a significant difference at P < 0.05.

### The effects of TEN on Heart rate variability and galvanic skin conductance during acute stress

In Experiment 3 (Fig. 2e) we next investigated the influence of TEN on sympathetic responses in a different group of volunteer subjects, who were challenged using a classical fear conditioning paradigm followed by a series of time-constrained cognitive tests to experimentally induce acute stress. We examined the influence of TEN on heart rate variability (HRV), a common biometric of psychophysiological arousal reflecting autonomic function including parasympathetic and sympathetic nervous system activity^39,40^. To experimentally increase sympathetic activity, we subjected volunteers to a stress trial in a between subjects design (sham and TEN n = 10 subjects per group) that included a classical fear-conditioning paradigm and time pressured cognitive tasks (Fig. 2e; see Methods). An independent t-test revealed there were no significant differences in average heart rate (HR) between the sham (HR = 67.01 ± 8.47 bpm) and TEN (HR = 69.18 ± 7.34 bpm) treatment groups during the stress trial (P = 0.55; Fig. 5a). Likewise the R-R intervals were not significantly different between the treatment groups during the stress paradigm (sham R-R interval = 922.07 ± 124.88 msec, TEN R-R interval = 885.21 ± 94.34 msec; P = 0.466; Fig. 5b). Analysis of HRV revealed subjects treated with TEN had a significant 34% reduction in SDNN compared to subjects treated with sham during the stress trial (sham SDNN = 105.45 ± 28.91 msec, TEN SDNN = 78.48 ± 16.60 msec; P = 0.02; Fig. 5c). Additionally, subjects in the TEN treatment group had a significant 88% reduction in the power of the LF (0.04 – 0.15 Hz) HRV component of the HRV spectrum compared to subjects treated with sham during the stress trial (sham LF power = 2946.80 ± 1352.12 msec^2^, TEN LF power = 1567.40 ± 827.24 msec^2^; P = 0.01; Fig. 5d). A similar trend was observed for the power of the HF component (0.15 – 0.4 Hz) of the HRV spectrum. Between group differences in the HF component of the HRV spectrum failed to reach a significant threshold (sham HF power = 2367.70 ± 1534.22 msec^2^, TEN HF power = 1381.50 ± 804.42 msec^2^; P = 0.09; Fig. 5f). There were no significant differences in the LF/HF ratios across treatment groups (sham LF/HF = 1.42 ± 0.43, TEN LF/HF = 1.20 ± 0.50; P = 0.30; Fig. 5f).

**Figure 5.**
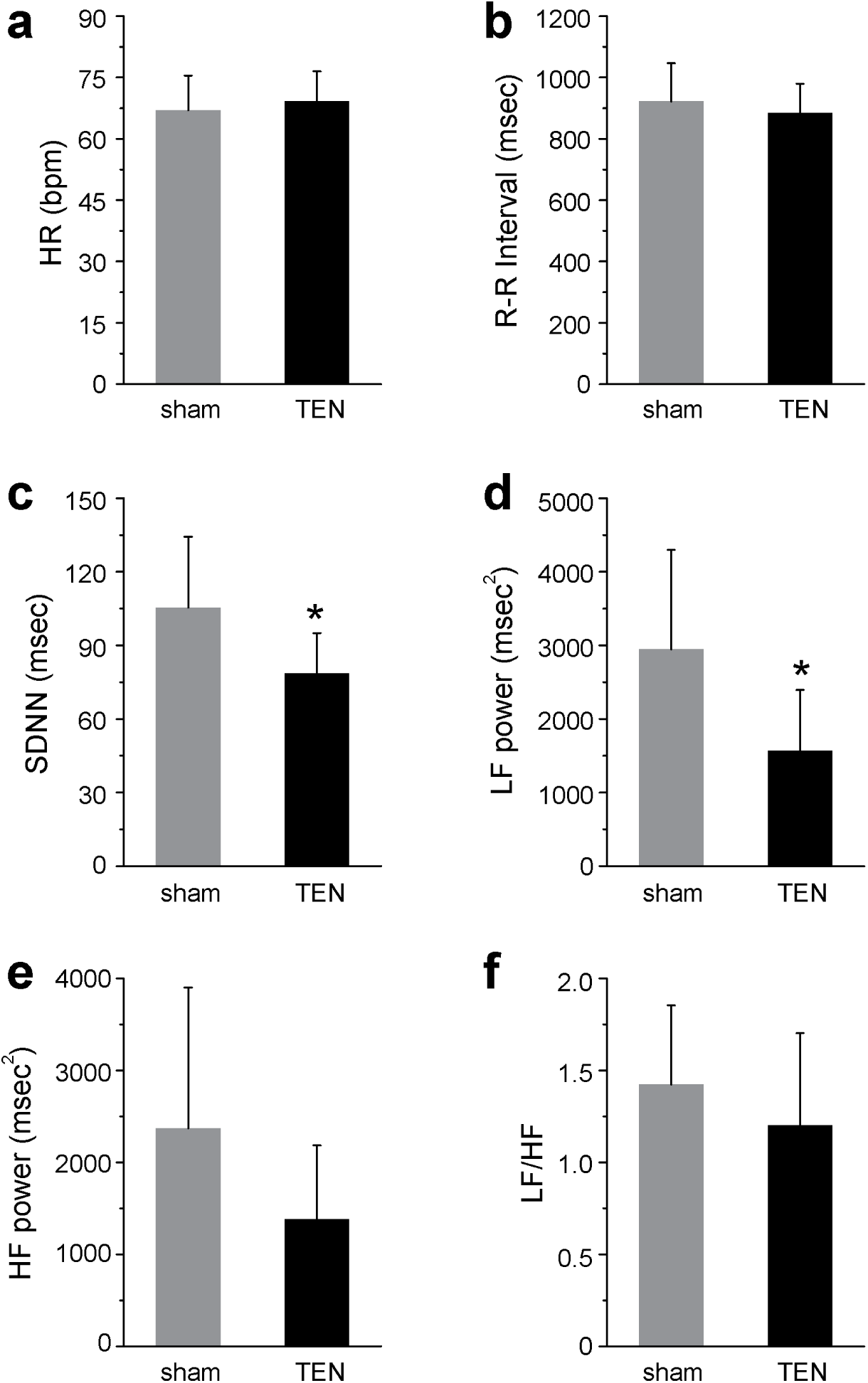
TEN suppresses changes in HRV induced by experimentally induced stress. The histograms illustrate the average HR (**a**), R-R interval (**b**), SDNN (**c**), LF power (**d**), HF power (**e**), and LF/HF (**f**) HRV responses mediated by sham and TEN treatments in response to the induction of acute stress. An asterisk indicates a significant difference with P < 0.05. All data shown are mean ± SD.

We also examined how TEN affected changes in electrodermal activity or galvanic skin conductance (GSC), which is another component of the SSR and is known to be sensitive to stress (Figs. 6a,b). Compared to sham we observed that TEN treatment (n = 10 subjects per group) produced a significant 32% suppression in anticipatory GSC change (ΔGSC) occurring in response to the onset of the fear-conditioning component of the stress trial prior to the delivery of the first unconditioned stimulus (ΔGSC_fear_; sham ΔGSC_fear_ = 0.54 ± 0.11, TEN ΔGSC_fear_ = 0.37 ± 0.14; P = 0.007; Fig. 6c). Similarly, we observed the average ΔGSC occurring in response to delivery of the unconditioned stimuli (10 electrical shocks per trial) was significantly suppressed by 53% in response to TEN treatment compared to sham treatment (sham ΔGSC_shock_ = 0.19 ± 0.04, TEN ΔGSC_shock_= 0.09 ± 0.06; P = 0.001; Fig. 6c). These data indicate TEN treatment significantly suppressed sympathetic activity in response to the delivery of unconditioned fear stimuli (electrical shocks) as indicated by the GSC component of the SSR.

**Figure 6.**
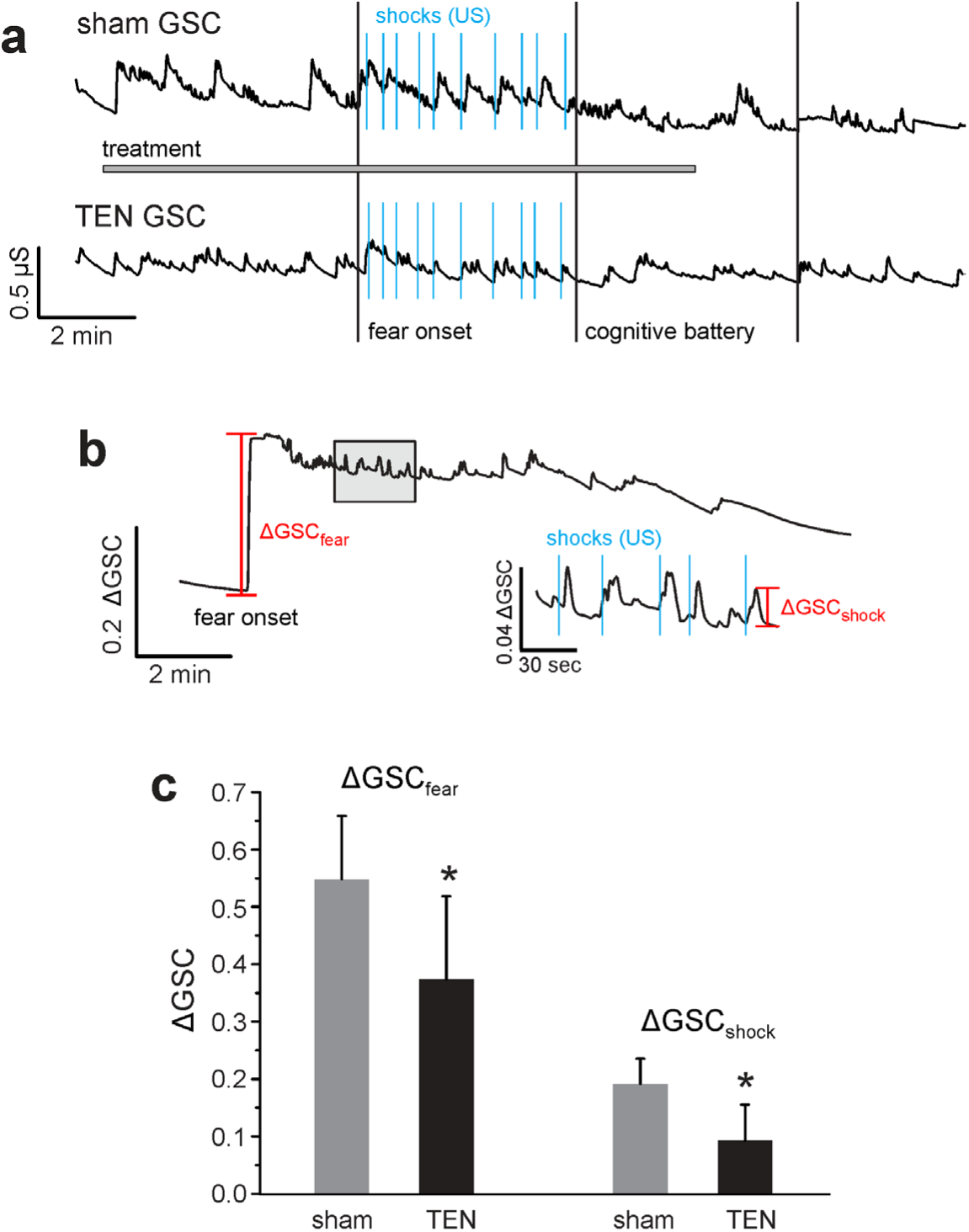
TEN attenuates anticipatory and event-related changes in GSC during classical fear-conditioning. **a**, Representative raw GSC traces obtained from a subjects treated with sham or TEN during the experimental paradigm designed to induce acute stress (Fig. 2e). The onset of the classical fear-conditioning paradigm (fear onset), delivery of unconditioned stimuli (US; electrical shocks; blue), and onset of the time-pressured cognitive battery are marked. Note differences in the amplitude of event-related GSC changes (ΔGSC; see Methods) between the treatment groups illustrate TEN-mediated reduction in sympathetic drive. **b**, The line plot illustrates a different GSC profile obtained from a subject treated with TEN during the fear-conditioning component of the stress trial. As also shown in panel **a**, all subjects exhibited the same type of response where there was a sharp anticipatory increase in GSC (ΔGSC_fear_) at the onset of the fear-conditioning component of the stress trial before the delivery of the first US. All subjects also exhibited transient GSC increases (ΔGSC_shock_) in response to the delivery of each US as illustrated. For clarity, the area marked by the *shaded rectangular box* is shown at a higher temporal resolution and amplitude scale (inset). **c**, The histograms illustrate the average ΔGSC for the TEN and sham treatment groups obtained during the fear-conditioning phase of the stress trial. An asterisk indicates a significant difference with P < 0.05. All data shown are mean ± SD.

### The influence of TEN on stress biomarkers: salivary α-amylase and cortisol

The protein enzyme α-amylase is widely recognized as a biochemical marker of sympathetic nervous system activity and sympathoadrenal medullary (SAM) axis activation^41-44^. More specifically, salivary levels of α-amylase directly correlate with plasma norepinephrine (NE) concentrations following the induction of acute stress including when electrical shock is used as a stressor^41-47^. To assess the impact of TEN on sympathetic activity and SAM axis activation, we examined fluctuations in salivary α-amylase (sAA) levels before (baseline) and after the acute induction of stress (Fig. 2e). We found that the mean baseline levels of sAA were not significantly different between the sham (n = 8 subject samples) and TEN (n = 10 subject samples) treatment groups (sham = 102.45 ± 39.60 U/mL, TEN = 85.57 ± 41.03 U/mL; P = 0.39; Fig. 7a). Ten minutes after the stress trial, the average levels of sAA for the sham group had increased 0.6% from their baseline values while in the TEN-treatment group they had dropped 24%. However, this was not a significant difference between treatment groups (sham = 103.13 ± 46.43 U/mL, TEN = 68.93 ± 27.26; P = 0.07; Fig. 7a). Thirty minutes after the stress trial, the average sAA levels for the sham group had increased 6.5% from their baseline values while the sAA levels for subjects treated with TEN had dropped 19.8% from baseline values resulting in a significant difference between treatment groups (sham = 110.30 ± 35.11 U/mL, TEN = 68.62 ± 28.93 U/mL; P = 0.01; Fig. 7a).

**Figure 7.**
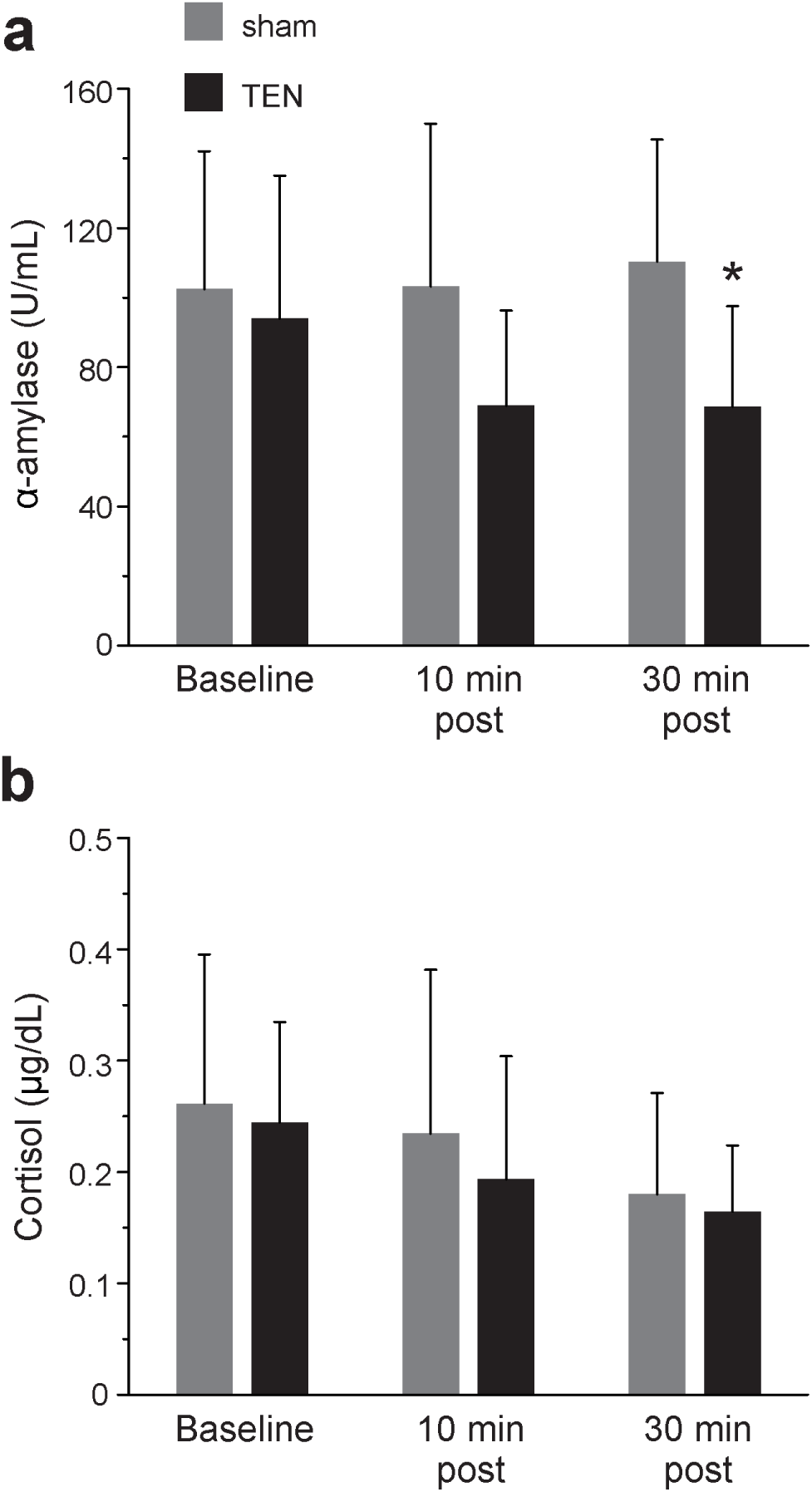
TEN reduces salivary α-amylase levels in response to acute stress. The histograms illustrate the average levels of α-amylase (**a**) and cortisol concentration (**b**) in saliva samples taken before (baseline), 10 min, and 30 min after the stress trial for sham and TEN treatment groups. An asterisk indicates a significant difference with P < 0.05. All data shown are mean ± SD.

While sAA is reflective of NE levels and acute SAM axis activity, cortisol is another stress biomarker, which has a slower onset, longer acting time course, and is under hormonal control of the hypothalamic-pituitary-adrenal (HPA) axis. It has previously been shown that sAA and cortisol can exhibit different response profiles that do not always correlate with one another depending upon stressor properties, such as the emotional intensity/valence of a stimulus and the duration of stress exposure^45-48^. Salivary cortisol levels did not significantly differ between sham and TEN treatments at any of the time points tested. At baseline the salivary cortisol concentration for sham was 0.26 ± 0.13 μg/dL and 0.24 ± 0.09 μg/dL for the TEN-treatment group (P = 0.75; Fig. 7b). Ten minutes after the stress paradigm the salivary cortisol concentration for sham was 0.23 ± 0.15 μg/dL and 0.19 ± 0.11 μg/dL for the TEN-treatment group (P = 0.51; Fig. 7b). Thirty minutes after the stress paradigm the salivary cortisol concentration for sham was 0.18 ± 0.09 μg/dL and 0.16 ± 0.06 μg/dL for the TEN-treatment group (P = 0.67; Fig. 7b).

### The effects of TEN on cognitive performance or executive processing reaction times

Subjects took three consecutive, time-constrained cognitive tests (Flanker test, n-back, and Stroop task; 2 min each) evaluating attention and working memory to evaluate the effects of TEN on executive processing during stress induction. A series of one-way ANOVAs revealed there were no significant differences between sham and TEN treated subjects in error rates or reaction times for congruent and incongruent tasks (Table 1). These data indicate that while TEN can suppress sympathetic activity in response to acute stress, it does so without impeding general/executive cognitive performance.

**Table 1.** Effects of TEN and sham treatment on cognitive performance.

|  | TES |  |  | Sham |  |  | Comparison |  |
| --- | --- | --- | --- | --- | --- | --- | --- | --- |
|  | n | M | SD | n | M | SD | ANOVA | P |
| **Flanker** |  |  |  |  |  |  |  |  |
| % Correct | 10 | 100.00 | 0.00 | 10 | 99.00 | 2.11 | F (1, 19) = 2.25 | 0.15 |
| Congruent RT | 10 | 612.37 | 249.27 | 10 | 570.43 | 190.43 | F (1, 19) = 0.18 | 0.68 |
| Incongruent RT | 10 | 585.92 | 170.93 | 10 | 618.00 | 182.06 | F (1, 19) = 0.17 | 0.69 |
| **Stroop** |  |  |  |  |  |  |  |  |
| % Correct | 8 | 98.75 | 3.54 | 6 | 97.82 | 2.79 | F (1, 13) = 0.28 | 0.61 |
| Congruent RT | 8 | 973.41 | 305.72 | 6 | 931.88 | 132.58 | F (1, 13) = 0.10 | 0.76 |
| Incongruent RT | 8 | 1247.82 | 289.62 | 6 | 1102.69 | 170.31 | F (1, 13) = 1.18 | 0.30 |
| **N-Back** |  |  |  |  |  |  |  |  |
| % Correct | 8 | 78.75 | 10.69 | 9 | 83.75 | 13.42 | F (1, 16) = 0.71 | 0.41 |
| Average RT | 8 | 705.80 | 103.74 | 9 | 713.25 | 133.01 | F (1, 16) = 0.02 | 0.90 |
| Combined Time | 8 | 921.80 | 245.53 | 9 | 890.29 | 291.78 | F (1, 16) = 0.06 | 0.81 |

M = mean, SD = standard deviation, RT = reaction time in milliseconds.

### TEN did not induce significant side effects

We assessed differences in the incidence (frequency), duration (in minutes), and severity (eight point scale) of the most common side effects associated with electrical neurostimulation using subject self-reports. TEN and sham groups had similar incidence rates of side effects with similar severity and durations (Table 2). Notably, there were no reports of headaches, nausea, or hearing changes. Further, no severe adverse events were reported.

**Table 2.** Summary of side effects elicited by TEN and sham treatments.

|  | TES |  |  |  |  |  | Sham Stimulation |  |  |  |  |  |
| --- | --- | --- | --- | --- | --- | --- | --- | --- | --- | --- | --- | --- |
|  | n | Presence | Severity |  | Duration |  | n | Presence | Severity |  | Duration |  |
|  |  | % (Count) | M | SD | M | SD |  | % (Count) | M | SD | M | SD |
| Skin Irritation | 9 | 33 (3) | 3.50 | 2.12 | 5.00 | 0 | 10 | 50 (5) | 4.20 | 2.49 | 2.65 | 4.90 |
| Headache | 10 | 0 (0) | -- | -- | -- | -- | 10 | 0 (0) | -- | -- | -- | -- |
| Dizziness | 10 | 0 (0) | -- | -- | -- | -- | 10 | 10 (1) | 3.00 | -- | 0.17 | -- |
| Vision Changes | 10 | 10 (1) | 3.00 | -- | 10.00 | -- | 10 | 0 (0) | -- | -- | -- | -- |
| Hearing Changes | 10 | 0 (0) | -- | -- | -- | -- | 10 | 0 (0) | -- | -- | -- | -- |
| Nausea | 10 | 0 (0) | -- | -- | -- | -- | 10 | 0 (0) | -- | -- | -- | -- |

M = mean, SD = standard deviation

## DISCUSSION

Evidence accumulated in Experiment 1 demonstrating TEN affected emotional thermoregulation by significantly increasing facial temperatures compared to sham is indicative of dampened basal sympathetic activity (Fig. 3b). The POMS data from Experiment 2 demonstrate that TEN significantly reduced anxiety and tension compared to sham treatment (Fig. 4). These observations are consistent with other reports investigating sympathetic skin responses (SSR) using functional infrared thermography that have shown increased sympathetic activity causes a decrease of facial temperatures while decreased sympathetic activity or relaxed mental states evoke increased thermal fluctuations of the face^30-37^. When we challenged subjects in Experiment 3 and induced acute stress using a classical fear-conditioning paradigm, we observed that TEN significantly suppressed stimulus-evoked galvanic skin responses, which represent another component of the SSR (Fig. 6c). In Experiment 3 we also observed that TEN produced a significant reduction in the levels of salivary α-amylase (sAA) compared to sham treatment (Fig. 7a). This particular observation suggests TEN suppressed sympathetic activity, in part, by modulating noradrenergic signaling pathways since previous studies have shown sAA levels to be predictive of plasma norepinephrine (NE) concentrations^44-50^. Taken together our observations indicate that TEN influences sympathetic activity, in part, by acting on noradrenergic signaling: TEN can dampen basal sympathetic tone, reduce tension and anxiety, and suppress psychophysiological and biochemical responses elicited by acute stress.

Based on anatomical, physiological, and biochemical observations reported here and elsewhere in the literature, we propose that TEN transmitted via cranial nerve and cervical spinal afferent pathways modulated the activity of the locus coeruleus (LC) and noradrenergic signaling to attenuate sympathetic activity and physiological stress. We developed the TEN method used in this study to target the ophthalmic and maxillary branches of the trigeminal nerve, the temporal branch of the facial nerve, and cervical spinal nerves (Fig. 2a). Trigeminal nuclei in the brain stem provide inputs to the LC and establish neural pathways by which trigeminal afferents can modulate noradrenergic activity (Fig. 1)^16^. Afferent projections from cervical spinal pathways provide another pathway by which TEN could have modulated the LC^17^. Afferent modulation of LC activity at a minimum would alter the mobilization and release of NE to mediate the brain’s response to environmental threats and stress^51^. It has indeed been shown that afferent sensory stimulation rapidly increases the firing rates of NE neurons in the LC^52,53^. In turn, modulation of NE signaling would have an impact on sympathoadrenal medullary (SAM) axis activity as further discussed below. In addition, pathways emerging from the primary sensory and spinal nuclei of the trigeminal nerve, sensory afferents from the facial nerve, project to the NTS, which also influences the activity of neurons in the RAS including the LC. The NTS also influences the activity of the dorsal motor nucleus (DMN) of the vagus, which sends preganglionic parasympathetic fibers to various target organs and regulates physiological homeostasis. At present, we cannot rule out the possibility that TEN may have also affected parasympathetic activity; however, our targeting approaches are anatomically and physiologically consistent with effects on sympathetic activity that involve impingement upon the LC.

We chose to implement high-frequency pulsing parameters for the development and implementation of TEN since it minimized the activation of neuromuscular and pain fibers. This is consistent with the observations of others showing high-frequency (10 kHz) spinal cord stimulation has been found to be effective at blocking low back and leg pain without activating nociceptive fibers or producing paresthesia^54^. Similarly high-frequency electrical stimulation (2 - 20 kHz) has been shown to be capable of rapidly and reversibly blocking neuromuscular activity^55^. One must also consider evidence dating back to the dawn of modern electrophysiology demonstrating that high-frequency currents can effectively stimulate peripheral nerves through a variety of mechanisms including by avoiding a refractory period^56^. The precise actions of TEN on the activity of specific nerve fiber types or afferent pathways is presently unknown. Future studies using electrophysiological approaches will need to be conducted to accurately characterize the influence of TEN on nerve fiber activity and downstream brain circuitry.

We hypothesize that through cranial and cervical spinal nerve afferent pathways TEN drives LC activity into a sympatholytic phasic firing mode by diminishing high levels of stress-induced tonic activity (Fig. 8b). The LC exhibits two essential types of neural discharges (tonic and phasic) that modulate cognition and behavior. These phasic and tonic LC discharge patterns favor different modes of signal processing and behaviors in a manner dependent upon the level of afferent drive^19,27,57^. In response to sensory and environmental cues, neurons in the LC have been shown to undergo shifts from a tonic to a phasic firing mode^58^. Based on experimental observations in animal models, the magnitude of endogenous phasic LC activity depends on the level of tonic LC activity (Fig. 8a). Under stress, tonic activity is high and phasic activity is diminished (Fig. 8a)^57,59,60^. Likewise when tonic activity is extremely low, such as during slow wave sleep, phasic activity is also diminished^57,59,60^. Using optogenetic control of mouse LC neurons *in vivo* it has recently been shown that tonic and phasic activity produce robust changes in sleep/wake cycles, motor behaviors, and general arousal in a frequency-dependent manner^61^. These observations reinforce the notion that appropriate levels of arousal stemming from a balance between tonic and phasic LC discharges and noradrenergic activity are necessary for maintaining optimal behavioral flexibility and performance^19,27,57^. Future investigations aimed at neurophysiologically elucidating the transfer function(s) between TEN stimulus parameters and LC tonic/phasic activity will be critical to gaining a complete understanding of how TEN influences sympathetic tone and stress responses.

**Figure 8.**
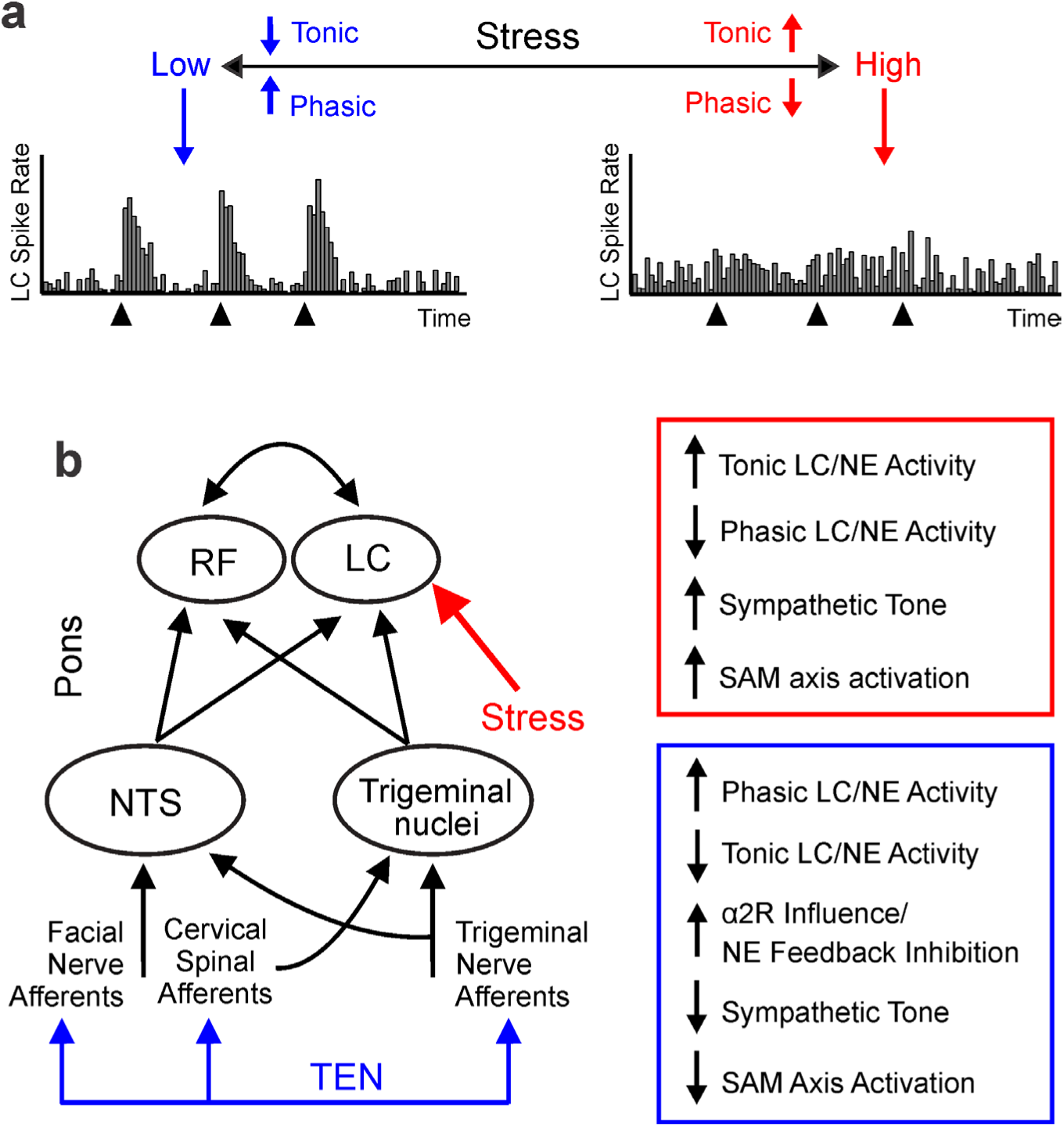
Neurons in the LC dynamically shift between tonic and phasic firing modes as a function of afferent activity and stress. **a**, The figure illustrates a working model of neuronal activity patterns exhibited by the LC under different levels of stress and arousal as observed from experimental animal models. This model incorporates observations that stress leads to high levels of tonic activity in the LC while sensory stimulation (black triangles) triggers phasic activity especially when tonic activity is low^52,53,57-60^. **b**, Facial and trigeminal nerve afferents, as well as the pons portion of the circuit diagram from Fig. 1 are shown (left). As illustrated stress (red) acts on LC neurons to increase their tonic firing rates thereby driving sympathetic activity and increasing SAM axis activation. We hypothesize that TEN (blue) biases LC activity through afferent pathways in a manner that increases phasic firing, reduces tonic activity, and suppresses sympathetic tone and SAM axis activation. Our observations suggest this partial mechanism of action, at a first station of information processing in the brain, may involve increased α2 adrenergic activity as depicted.

Our physiological and biochemical observations indicate TEN may partially act by producing an increase in α2 adrenergic receptor activity to inhibit NE levels through known negative feedback mechanisms. Clonidine, an α2 adrenoceptor agonist, has been shown to block sAA secretion triggered by sympathetic fiber stimulation^62^. Similarly, we observed that TEN significantly attenuated the levels of sAA following stress induction as discussed above (Fig. 7). Interestingly, clonidine inhibits the tonic activity of LC neurons without suppressing phasic activity^63^. Clonidine has also been shown to exert some of its actions by significantly decreasing the LF component of HRV while producing minimal effects on the HF component of HRV in humans^64^. Others have also reported the oral administration of clonidine to healthy subjects significantly reduces the LF component of HRV, as well as lowers plasma concentrations of NE^65,66^. Similar to these actions of clonidine, we found that TEN treatments produced a significant decrease in the LF component of HRV during the stress trial compared to sham (Fig. 5d,e). Further experiments involving human pharmacology will be required to fully delineate the molecular mechanisms of action underlying the ability of TEN to inhibit sympathetic activity. These will be important studies given the growing interest of utilizing peripheral nerve pathways to affect brain function, such as efforts exploring the development of electroceuticals^67^.

The safety of TEN is supported by a long history of peripheral electrical neurostimulation obtained over more than four decades. Legally marketed electrical nerve stimulation devices are already commercially available and have output levels in excess of 100 mA, which is far greater than current amplitudes implemented in this study. Some of these devices are intended for over-the-counter cosmetic applications of transcutaneous electrical nerve stimulation (TENS) and affect the maxillary branch of the trigeminal nerve, which has some overlap with neural pathways we targeted. One example device is the Bio-medical Research (BMR) Face device, which is an over-the-counter TENS device designed to target the trigeminal nerve and provide neuromuscular electrical stimulation (NMES) waveforms to encourage facial rejuvenation for aesthetic purposes. A recent study examined the safety and efficacy of this device at a peak current intensity of 35 mA when used five days per week for 20 minutes each day for 12 weeks^68^. There were no significant adverse events in this study and the only reported side effects were minor skin redness following stimulation, which disappeared with 10-20 minutes following use^68^. Consistent with these results, on an acute level we found that TEN did not elicit any significant side effects (Table 2). Further, TEN did not have any significant effect on cognitive performance or reaction times (Table 1). Other observations demonstrating the chronic safety profile of TEN have been recently reported^69^. Based collectively on these observations, we conclude TEN provides low-risk and reliable approach to modulating psychophysiological arousal.

In summary our observations demonstrate that, compared to sham, TEN significantly dampened basal sympathetic tone, significantly decreased tension and anxiety, and significantly suppressed the physiological and biochemical responses to acute stress induction. The reduction of LF HRV, decreased levels of sAA, and suppressed GSR in response to acute stress are consistent with the hypothesis that TEN modulation of trigeminal (V1/V2) and cervical spinal (C1/C2) afferent pathways act partially through bottom-up mechanisms to modulate noradrenergic activity in a first station of information integration in the human brain. While further experiments including electrophysiological assays and pharmacological manipulations will be required to test this hypothesis, as well as to investigate the influence of TEN on signaling cascades downstream of the LC and NE, our basic observations have provocative implications for the management of stress and optimization of brain health.

## METHODS

### Participants

All experimental procedures were conducted on human volunteers in accordance with protocols guidelines approved by an Institutional Review Board (Solutions IRB, Little Rock, AR). All subjects provided written informed consent prior to experimentation. We conducted three independent experiments in this study using different groups of volunteer subjects. All experiments were designed using a between subjects approach. All subjects were blinded to the study conditions. For all experimental conditions, exclusion criteria were as follows: neurological or psychiatric disorder, cranial or facial metal plate or screw implants, severe face or head trauma, recent concussion or brain injury, recent hospitalized for surgery/illness, high blood pressure, heart disease, diabetes, acute eczema on the scalp, and uncorrectable vision or hearing.

Experiment 1 (Fig. 2c) was designed to study the influence of TEN on a component of the sympathetic skin response by implementing functional infrared thermography to assay changes in emotional thermoregulation expressed in several facial regions as described below. The sample pool was consisted of 19 healthy right-handed subjects (7 male, 12 female) between the ages of 18 to 27 (mean age = 22.16 ± 2.09 years). Subjects were randomly assigned to receive sham (n = 10) or TEN (n = 9). The ethnicity of the subjects was as follows: 42% of participants were Asian, 37% were Caucasian, 10% were African-American, and 5% were Hispanic. The educational background of subjects was as follows: 47% had completed some college, 31% had a Bachelor’s degree, and 20% had completed some post-graduate work or had a post-graduate degree.

Experiment 2 (Fig. 2d) was designed to test the impact of TEN on affective mood as reported by the Profile of Mood States (POMS) survey^38^. The sample pool consisted of 45 healthy subjects (16 male, 29 female) that ranged in age from 18 to 43 (mean age = 22.46 ± 4.23 years). Subjects were randomly assigned to receive sham (n = 20) or TEN (n = 25). The ethnicity of subjects was as follows: 47% were Caucasian, 31% were Asian, 16% were African-American, and 4% were Hispanic. The educational background of subjects was as follows: 44% completed some college, 29% had a Bachelor’s degree, and 16% completed some post-graduate work or had a post-graduate degree.

Experiment 3 (Fig. 2e) was designed to study the influence of TEN on psychophysiological arousal and the mobilization salivary biochemicals in response to acute stress induced by a classical fear conditioning paradigm and a series of time pressured cognitive tasks. The sample pool consisted of 20 male subjects (sham n = 10; TEN n = 10) to avoid the introduction of confounds related to hormonal variance across menstrual cycles on stress biochemical profiles. The subjects were between the ages 19 to 27 (mean age = 22.3 ± 2.2 years). The ethnicity of subjects was as follows: 50% of the subjects were Asian, 35% were Caucasian, 10% were African American and 5% were Hispanic. The educational background of subjects was as follows: 40% of participants had completed some college, 40% had a Bachelor’s degree, and 15% had completed some post-graduate work.

### Transdermal electrical neuromodulation

Prior to this study, we spent two years developing and investigating a variety of electrical neuromodulation waveforms and approaches. The transdermal electrical neuromodulation (TEN) waveform developed for use in this study was a pulsed (7 – 11 kHz; 50% duty cycle) biphasic current producing average amplitude of 5 – 7 mA for 15 min (Experiments 1 and 2; Figs. 2c,d) or 14 min (Experiment 3; Fig. 2e). The TEN waveform pulse frequency linearly ramped from 7 – 11 kHz during the first 30 sec of the protocol and ramped down linearly from 11 – 7 kHz during the last 30 sec of the stimulus. The sham waveform was an active stimulation control as described above, but pulsed at a frequency of 1 - 2 kHz (50% duty cycle; < 4 mA average current amplitude) for 15 min (Experiments 1 and 2; Figs. 2c,d) or 14 min (Experiment 3; Fig. 2e) to mimic skin sensations similar to those experienced throughout the real TEN stimulation protocol (Table 2). Subjects were only able to control the stimulus intensity within the ranges indicated, but not the frequency of stimuli. Subjects were not able to distinguish any differences between the sensations elicited by the real TEN or sham waveforms. During both the real TEN and sham stimulus protocols, subjects were instructed to adjust the current output of a medical-grade wearable TEN device (Thync, Inc., Los Gatos, CA) using an iPod touch connected to the device over a Bluetooth low energy network such that it was comfortable. TEN and sham waveforms were delivered to the right temple (10/20 site F8) and base of the neck (5 cm below the inion) using custom-designed electrodes comprising a hydrogel material and a conductive Ag/AgCl film secured to the wearable TEN device. The anterior electrode positioned over F8 was a 4.9 cm^2^ oval having a major axis of 2.75 cm and a minor axis of 2.25 cm while the posterior electrode was a 12.5 cm^2^ rectangle with a length of 5 cm and a height of 2.5 cm. The average current density was < 2 mA/cm^2^ at all times to keep in accordance with general safety practices to prevent any damage to the skin. Subjects were assigned to experimental conditions using a randomization method or a counterbalancing approach. Subjects were always kept blind to all experimental conditions.

### Functional infrared thermography

In Experiment 1 (Fig. 2c), after providing brief demographic information, subjects were seated in front of a calibrated infrared thermal imaging camera (Fig. 2a; FLIR T450sc, FLIR Systems Inc., Nashua, NH) positioned 1.5 meters from subjects face in a thermally stable testing room (Fig. 2b) maintained at 24 °C. Time-lapsed (30 Hz) infrared (λ = 7.5 to 13 μm) images were acquired during the 5 min baseline period, during the 15 min sham or TEN treatment period, and for a recovery period up to 10 min following the termination of treatment (Supplementary Videos 1 and 2). Images were then stored for later offline image analysis. Regions of interest were positioned and tracked on the forehead, cheeks, nose, and chin region of the face (Fig. S1). From radiographic datasets the average baseline temperature was calculated for each subject and facial location across the 5 min baseline period. Average temperatures were calculated for each subject from each facial region during the 2, 5, 10, and 20 min period following the onset of sham or TEN stimulation (Fig. 3a). These average temperatures were analyzed and expressed as a percent change from baseline (Fig. 3b) for each subject to account for any variation in baseline temperatures across subjects.

### Profile of Mood States Survey

In Experiment 2 (Fig. 2d), after providing brief demographic information, subjects were seated in a testing room (Fig. 2b). Subjects had a 5 min resting period prior to receiving a 15 min sham or TEN treatment. Immediately following stimulation, participants completed the POMS survey^38^, which is a 65-item scale designed to assess transient affective states and comprises six subscales: Anger-Hostility (11 items), Depression-Dejection (14 items), Fatigue-Inertia (8 items), Vigor-Activity (6 items), Tension-Anxiety (3 items), and Confusion-Bewilderment (3 items). Items were scored on a 5-point scale, ranging from 0 = not at all to 4 = extremely and were indexed to how participants felt in that moment. Reliabilities of the subscales ranged from 0.50 to 0.91.

### Acute stress induction

In Experiment 3 (Fig. 2e) we implemented an acute stress induction paradigm to study the effects of TEN on physiological and biochemical stress responses. Subjects received either TEN or sham treatment in a between subjects design (N = 10 subjects per group). All participants were tested between the hours of 13:00 and 16:00 to limit variability introduced by circadian fluctuations in salivary analytes (*see Salivary collection and stress biomarker assays below*). Following informed consent, subjects were allowed to acclimate for 20 min before providing a baseline saliva sample. After providing the initial saliva sample, subjects were connected to a wearable TEN device as described above, a peripheral nerve stimulator (MiniStim MS-IVA, Life-Tech, Inc., Dallas, TX) was positioned over the median nerve of the right wrist, and a Shimmer3 (Shimmer, Dublin, Ireland) optical heart rate (HR) monitor and galvanic skin conductance (GSC) sensor was placed on the index, middle, and ring fingers of the opposite hand. The timing, presentation of stimuli, and acquisition of HR and GSC data was accomplished using Attention Tool (iMotions, Inc., Cambridge, MA).

The stress trial (Fig. 2e) commenced 30 sec prior to the onset of TEN or sham treatment. On the beginning of the stress trial there was a 6 min pre-trial period to give the TEN or sham treatment time to begin exerting an effect before the induction of acute stress began. The stress trial comprised a 6 min classical fear-conditioning paradigm immediately followed by a 6 min time-constrained cognitive battery. Immediately following the time pressured cognitive tests the stress trial concluded with a 3 min neutral video of a nature scene. Both the classical fear-conditioning paradigm and the time-constrained series of cognitive tests are known to induce acute stress and increase sympathetic activity^70-73^. Before the beginning of the stress trial, participants were instructed that when the computer monitor they were seated in front of began to flash still images (instead of the baseline period video) that they would be given an electrical shock every time an image of lightning appeared. The transition from baseline videos to still images during the fear-conditioning portion of the stress trail induced an anticipatory increase in acute stress as reflected by instantaneous changes in GSC (Fig. 6b) in every subject. During the fear-conditioning component of the stress trial participants were randomly presented with 40 still images of nature scenes for 6 seconds each: 10 images of lightning paired with electrical shock (0.5 sec, 4 - 6.5 mA) and 30 neutral nature scenes.

The second component of the stress trial included a time-constrained series of three cognitive tests (2 min each) including a Flanker test, n-back working-memory test, and a Stroop task. The Flanker task is a selective attention task in which participants indicate the direction of a target stimulus that is flanked by stimuli that are oriented in the same direction (congruent), in the opposite direction (incongruent), or a neutral direction as the response target. The n-back, which assesses working memory, has participants view a sequence of stimuli and indicate when the current stimulus matches the stimuli from n steps earlier in the sequence. In this case, subjects were instructed to get to 2 back as quickly and accurately as possible. The Stroop task tests sematic memory by having participants indicate the color of the ink in which a color word is written as fast as possible. Trials can be congruent, the text color and the word refer to the same color, or incongruent, the ink color and the word refer to different colors, which can lead to frustration and itself induce acute stress ^72,73^. Reaction times and accuracy were measured for all tests and analyzed off-line in subsequent analyses. Following the stress trial, there was a 30 min. recovery period during which subjects reported the presence, duration (in minutes) and severity (0 to 8 scale) of skin redness or irritation, headache, dizziness, nausea, and vision or hearing changes and two additional saliva samples were collected as described below.

### Heart rate variability and galvanic skin conductance metrics

We acquired cardiac activity and electrodermal activity during the stress trial using a Shimmer3 optical heart rate monitor integrated with a GSC sensor. Raw electrocardiogram data were collected using Attention Tool before being processed offline using Kubios HRV (University of Eastern Finland, Kuopio, Finland). From these data we quantified the average HR, R-R interval, standard deviation of the normal-to-normal heartbeat (SDNN), power in the low-frequency (0.04 – 0.15 Hz) and high-frequency (0.15 – 0.4 Hz) bands of the HRV spectra, and the LF/HF ratio in response to TEN and sham treatments during the stress trial. All GSC data were acquired using Attention Tool and stored offline for analysis using Igor Pro (Wavemetrics, Inc., Portland, OR). We quantified the peak-to-peak changes for GSC on raw data, where: ΔGSC = (GSC_peak_ – GSC_base_)/GSC_base_ occurring at the onset of the fear-conditioning component of the stress trial (ΔGSC_fear_; Fig. 6b) and the average peak-to-peak change in response to the delivery of the 10 unconditioned stimuli or electrical shocks (ΔGSC_shock_; Fig. 6b) for each subject.

### Salivary collection and stress biomarker assays

Prior to arrival, participants were instructed to not brush their teeth within 45 minutes, eat within one hour, consume caffeine or alcohol within 12 hours or have dental work performed within 24 hours of their scheduled appointment. After providing informed consent, participants rinsed their mouths in preparation for contributing a saliva assay and were then seated in a quiet room during which they self-reported basic demographic information. After 20 minutes, participants provided a baseline saliva sample via the passive drool method. As per manufacturer’s instructions (SalivaBio, Inc., State College, PA), saliva is pooled at the front of the mouth and eased through a tube, centered on the lips, directly into a cryovial, and immediately stored at −20° C. The same collection procedure was used to collect additional saliva samples 10 min and 30 min following the end of the stress trial. Saliva samples were coded and sent to Salimetrics, LLC (State College, PA) where ELISA methods were employed to assess α-amylase (Salimetrics 1-1902) and cortisol levels (Salimetrics 3002) in a blinded manner using a subject coding procedure.

The protein α-amylase is widely recognized as a biochemical marker of sympathetic nervous system activity and sympathoadrenal medullary (SAM) axis activation. More specifically, salivary levels of α-amylase directly correlate with plasma norepinephrine (NE) concentrations following the induction of acute stress including when electrical shock is used as a stressor^41-47^. Cortisol is a prototypical stress hormone reflective of hypothalamus pituitary adrenal (HPA) axis activation, which has slower onset and longer lasting effects compared to SAM axis activation.

### Statistical analyses

All statistical analyses were conducted using one-way ANOVA’s, repeated measures ANOVA’s, or independent t-tests with IBM SPSS Statistics Software (IBM Corporation, Armonk, NY) as indicated. Experimenters analyzing data were blinded to treatment groups using a subject coding procedure. Prior to analysis all group data were confirmed to be normally distributed using a test for normality with the Shapiro-Wilk procedure (P > 0.05) and visual inspection of Q-Q plots in SPSS. The variance on all between group data was examined using Levene’s test for homoscedasticity with no significant differences at P > 0.05. Analyses of thermographic data from different facial regions were conducted using a repeated measures ANOVA with time as the within subjects factor and treatment as the between subjects factor. The POMS data and cognitive data were analyzed using a series of one-way ANOVA’s with SPSS. The HR, HRV, and GSC data were analyzed using independent t-tests. Thresholds for statistical significance were set at P < 0.05. All data reported and shown are mean ± SD unless otherwise indicated.

## AUTHOR CONTRIBUTIONS

W.J.T., S.K.P., A.M.B. designed the experiments. W.J.T., A.M.B., H.M.M., R.S.S., J.D.C, M.A.M., K.A., L.A., D.Z.W, and S.K.P. conducted the experiments. W.J.T., A.M.B., H.M., R.S., J.D.C, M.A.M., K.A., L.A., D.Z.W, and S.K.P. analyzed data and assisted in the preparation of figures. W.J.T., A.M.B., J.D.C, M.A.M., K.A., L.A., D.Z.W, and S.K.P. assisted in the preparation and editing of the manuscript.

## ACKNOWLEDGMENTS

We thank Isy Goldwasser for insightful comments and critical feedback throughout our R&D efforts and in preparing this manuscript. We thank Wing Law, Ph.D., Jay Hamlin and Anil Thakur for their technical support in making this study possible. We are grateful to all the members of the Thync team for immense support and assistance throughout various aspects of platform development, device engineering, and system implementation.

## SUPPLEMENTARY INFORMATION

**Supplementary Figure 1.**
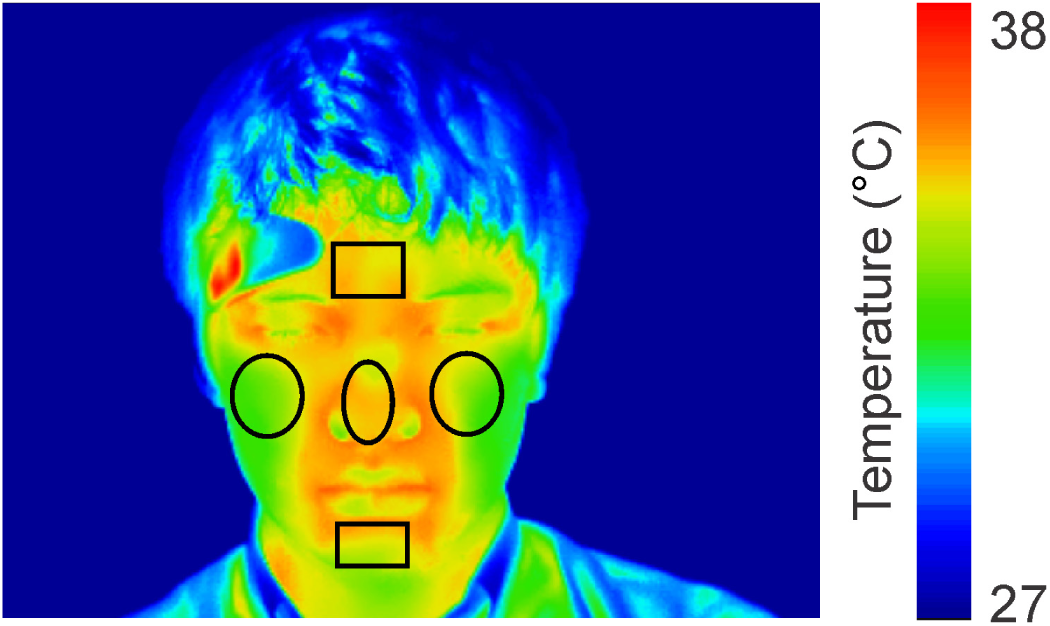
Approach to analysis of functional infrared thermography data. **a**, The image of radiographic thermal data shows regions of interest (black boxes) on the forehead, nose, cheeks, and chin that were used to measure the skin temperature at different time points from the time-lapsed data acquired. The image is shown with a pseudo-color look-up table applied to make the visualization of temperature differences easier for readers such that *blue* represents 27° C, *green* represents 33° C, and *red* represents 38° C.

